# An interoperable research agent network for scientific discovery

**DOI:** 10.64898/2026.08.13.744542

**Authors:** Tin Long Cheong, Xinchen Ji, Ying Wang, Yue Zhou, Bo Li, Cuiting Zhang, Jiabao Huang, I Wu, Ao Li, Edwin Cheung

## Abstract

Recent advances in agentic systems have enabled the autonomous execution of research tasks across scientific domains. However, the rapid emergence of specialized scientific agents for areas such as computational pathology, microbiome research, gene editing, materials science, organic chemistry, and drug discovery has created a fragmented ecosystem of scientific capabilities. While these agents often demonstrate strong performance within their respective domains, limited interoperability makes it difficult to combine expertise across platforms and coordinate complex interdisciplinary workflows. Here we introduce GUIA (Guided-research Utilizing Intelligent Agents), an interoperable research-agent network built upon a flexible Agent-to-Agent (A2A) communication architecture. GUIA enables both in-house and third-party agents to collaborate within shared workflows, allowing scientific capabilities to accumulate through the integration of complementary expertise. We evaluated GUIA through four assessments spanning baseline benchmarking, third-party single-agent integration, third-party multi-agent integration, and cross-server agent collaboration. Furthermore, we demonstrate its practical utility through real-world applications involving therapeutic target discovery, drug discovery, and spatial proteomics analysis. Together, our results show that interoperable research-agent networks can coordinate specialized expertise across independently developed systems, providing a scalable framework for expanding scientific capabilities through collaboration.

## Introduction

The growing complexity of scientific research increasingly demands interdisciplinary and cross-laboratory collaboration among experts. Such collaborations have often led to major scientific advances. For example, the Human Genome Project united hundreds of scientists across 20 institutions from six countries in a coordinated effort to sequence and analyse the human genome^1^. However, fostering effective collaboration requires substantial coordination among researchers with different expertise, as well as efficient sharing of knowledge, data, and results across laboratories. These requirements are often difficult to achieve in practice, creating barriers to addressing increasingly complex scientific questions.

Recent advances in large language models (LLMs) have created new opportunities for supporting interdisciplinary research. Trained on vast amounts of scientific literature and general knowledge, LLMs were initially applied to tasks such as literature summarization, scientific question answering, and facilitating scientific discussions^2,3^. More recently, LLMs have been coupled with external tools, databases, and computational environments, enabling them to autonomously execute increasingly sophisticated scientific workflows^4–8^.

Augmenting LLMs with computational tools has led to the rapid development of scientific agents across various research domains, including bioinformatics, computational pathology, gene editing, microbiome research, drug discovery, organic chemistry, and materials science^9–13^. For example, Pathology-CoT enables whole-slide image (WSI) analysis through pathology-specific reasoning strategies^9^, CRISPR-GPT automates gene editing design through integration of domain expertise and specialized tools^10^, and Eubiota coordinates multiple agents to support end-to-end microbiome research^11^. Similarly, ChemCrow augments LLMs with chemistry-specific tools for organic synthesis and materials design tasks^12^. Collectively these systems illustrate the growing diversity and specialization of scientific AI agents.

However, the rapid emergence of specialized agents has also created a fragmented ecosystem of scientific capabilities. Most agents are developed as independent systems optimized for specific domains, tasks, and computational environments. Consequently, the expertise embedded within one agent is often inaccessible to others. As the number of scientific agents grows, the lack of interoperability presents a significant barrier to coordinating expertise across domains and scaling AI-assisted scientific discovery. This limitation becomes particularly prominent in interdisciplinary research, where findings generated by agents in one system may need to be interpreted, validated, or extended by agents hosted in another. Similar challenges arise in cross-laboratory collaborations, where agents deployed on different servers may need to collaborate on shared scientific problems.

Consequently, the rapid emergence of specialized agents creates a growing need for enhanced interoperability. In parallel to efforts at developing increasingly capable scientific agents, a research-agent network offers a complementary strategy for combining expertise across independently developed systems. By enabling specialized agents to contribute their strengths within shared workflows, such networks allow scientific capabilities to accumulate through collaboration, providing researchers with access to expertise that spans multiple domains and computational environments.

To address this challenge, we introduce GUIA, an interoperable research-agent network built upon a scalable Agent-to-Agent (A2A) communication architecture^14^. GUIA enables human researchers to coordinate a network of specialized agents for complex interdisciplinary research tasks. The GUIA architecture is designed to support the formation and continual expansion of research-agent networks through the seamless integration of both in-house and third-party agents, irrespective of their underlying models, frameworks, tools, or deployment environment. By transforming a fragmented ecosystem of scientific agents into a unified network, GUIA facilitates the coordinated use of complementary expertise across diverse research domains. Furthermore, GUIA introduces a paradigm for cross-server collaboration, allowing agents hosted on different geographical infrastructures to coordinate on shared research tasks.

To evaluate GUIA, we conducted benchmark studies spanning multiple research domains and assessed its extensibility through the integration of both third-party single-agent and multi-agent systems. These integrations enabled GUIA to acquire capabilities beyond those available in its original configuration, demonstrating the extensibility of the research-agent network paradigm. We also evaluated the practical utility of GUIA through three real-world case studies: (1) designing and executing a customized drug discovery workflow; (2) analyzing spatial proteomics data to uncover novel biological patterns; and (3) identifying novel therapeutic targets. Together, these studies demonstrate how interoperable research-agent networks can coordinate specialized expertise across independently developed systems, providing a scalable foundation for AI-assisted scientific research.

## Results

### A scalable research-agent network for cross-disciplinary research

GUIA is a scalable framework of research agents built upon an Agent-to-Agent (A2A) communication architecture^14^. Under this framework, specialized agents can communicate and collaborate seamlessly, fostering an agent network for executing scientific tasks.

GUIA comprises a multi-layered architecture consisting of standalone specialized agents, each equipped with domain-specific tools and configurations (Fig. 1a). These agents are exposed through a standardized A2A protocol, enabling collaboration without the need for tight system integration (Fig. 1a). This approach allows complex research workflows to be decomposed into sub-tasks and delegated to the most relevant agents in an interoperable manner.

**Figure 1.**
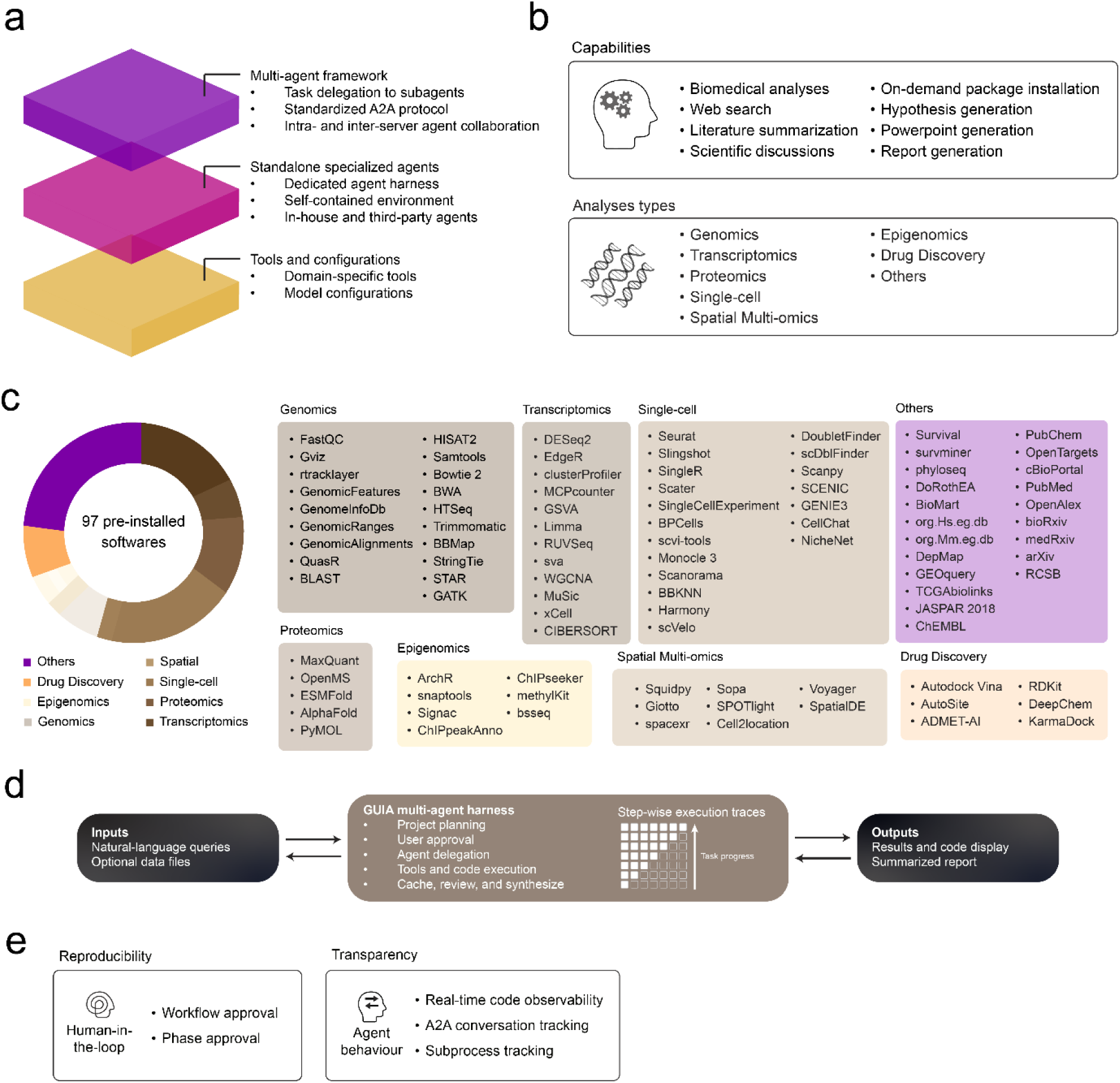
Architecture of GUIA. **a.** Overview of the multilayered architecture of GUIA. **b.** Capabilities and analyses supported by in-house agents. **c.** A unified execution environment exposing in-house agents to 97 tools. **d.** The human-AI working logic of GUIA. **e.** Multiple HITL and agent traceability features incorporated into GUIA.

Importantly, GUIA supports the collaboration of both in-house and third-party specialized agents. This capability enables agents built using different frameworks, large language models (LLMs), tools, and deployment environments to coordinate seamlessly (Fig. 1a). Our in-house agents are equipped with a robust execution environment that supports a broad range of research capabilities, including autonomous execution of biomedical analyses, on-demand installation of missing R and Python packages, web-based information retrieval, literature review, scientific discussion, hypothesis generation, and the automated creation of project reports and presentation materials (Fig. 1b and Fig. 2a). These agents have access to a total of 97 tools spanning multiple disciplines including genomics, transcriptomics, proteomics, single-cell biology, spatial multi-omics, epigenomics, drug discovery, and other research domains (Fig. 1c). Collectively, our in-house agents form the foundation of GUIA, upon which a growing network of third-party scientific agents can be incorporated to extend its capabilities across diverse research domains.

**Figure 2.**
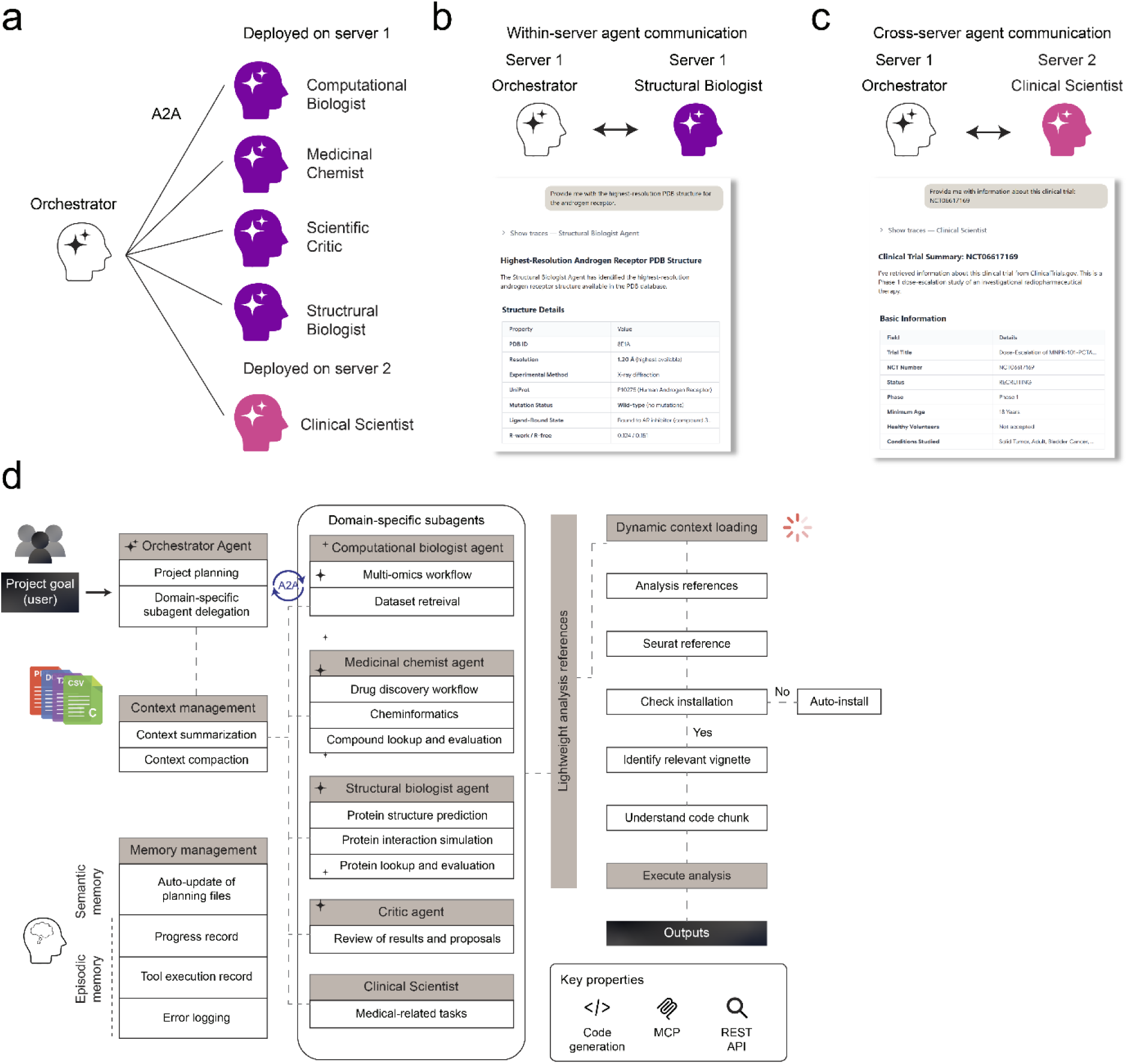
Characteristics of the multi-agent framework. **a.** Agent communication framework employed in GUIA. **b.** Example of a task delegated to the Structural Biologist agent hosted on server 1. **c.** Example of a task delegated to the Clinical Scientist agent hosted on server 2. **d.** Workflow of GUIA’s in-house agent architecture.

To facilitate accessibility, GUIA operates through a user-friendly natural language interface that allows researchers to submit research queries alongside optional data files (Fig. 1d). Upon receiving a request, GUIA evaluates the complexity of the task, presents a proposed workflow when appropriate, and coordinates scientific agents to execute the research plan (Fig. 1d). We further incorporated multiple human-in-the-loop (HITL) mechanisms to ensure user control and traceability of agent activities (Fig. 1e). Notably, all research outputs, together with agent reasoning, A2A conversation histories, tool invocations, and generated code, are recorded as traceable sub-processes, and displayed through a dedicated frontend interface (Supplementary Fig. 1).

### Characteristics of the research-agent network

GUIA employs a centralized architecture where an Orchestrator agent coordinates a collection of specialized agents (Fig. 2a). The Orchestrator is responsible for managing research projects, decomposing objectives into sub-tasks, and delegating them to domain-specific subagents. To demonstrate GUIA’s ability to coordinate a network of scientific agents, we deployed our in-house agents, including a Computational Biologist, Medicinal Chemist, Scientific Critic, Structural Biologist, and a Clinical Scientist across two geographically separate servers (Fig. 2a). Notably, each agent is exposed through an independent HTTP endpoint, enabling A2A communication via JSON-formatted responses^14^. Through two demonstrations, the Orchestrator effectively identified the required expertise for each task and delegated them to the most relevant scientific agents for execution. For example, a request to identify and retrieve structural information for the androgen receptor (AR) from the Protein Data Bank (PDB)^15^ was assigned to the Structural Biologist, which was hosted on the same server as the Orchestrator (Fig. 2b). On the other hand, a request to summarize information from a specific clinical trial was delegated to the Clinical Scientist, which was hosted on a geographically separate server (Fig. 2c). This highlights a paradigm for cross-server agent collaboration, in which AI agents developed by different institutions or hosted on different infrastructures can collaborate on shared scientific tasks.

Briefly, we adopted a modern agent-engineering framework for our in-house agents, whereby workflow references and project artifacts are dynamically loaded on demand during execution (Fig. 2d). This approach effectively reduces context bloat while steering agent actions toward best-practice research behaviours. To support long-horizon tasks, context volume is continuously monitored and, upon a threshold, compressed through automated summarization and compaction. In addition, GUIA maintains both semantic and episodic memory through the autonomous generation and update of planning files, tool execution records, and structured error logging, providing persistent project memory for both agents and the human researcher to revisit. We further leveraged a combination of agent engineering paradigms, including vibe coding^16^, the Model Context Protocol (MCP)^17^, and Representational State Transfer Application Programming Interfaces (REST-APIs) to facilitate code generation, tool deployment, information retrieval, and access to external databases (Fig. 2d).

### GUIA supports research capability expansion through third-party agent integration

GUIA is designed to transform isolated scientific agents into collaborative networks for tackling complex research tasks. We evaluated GUIA’s capabilities through a four-step assessment strategy. First, we evaluated its baseline performance on a set of biomedical research tasks. Specifically, GUIA was assessed on GenomeArena, a benchmark comprising 720 queries spanning seven domains: (1) drug–target relationships; (2) gene annotation; (3) gene–phenotype associations; (4) miRNA targets; (5) pathways; (6) protein structure; and (7) variant annotation^18^. Given that these tasks closely align with the expertise of GUIA’s in-house agents, they provide a measure of the system’s baseline capabilities. Across the seven evaluated domains, GUIA (GLM-5) achieved an average accuracy of 78%, surpassing base LLM (GLM-5; 56%) and Open Science^8^ (GLM-5; 71%), a multi-agent system composed of a fixed set of specialized agents (Fig. 3a, b). While GUIA demonstrated strong baseline performance through its in-house agents, its flexible architecture supports the integration of third-party scientific agents, allowing expertise from other systems to be combined within a unified research ecosystem.

**Figure 3.**
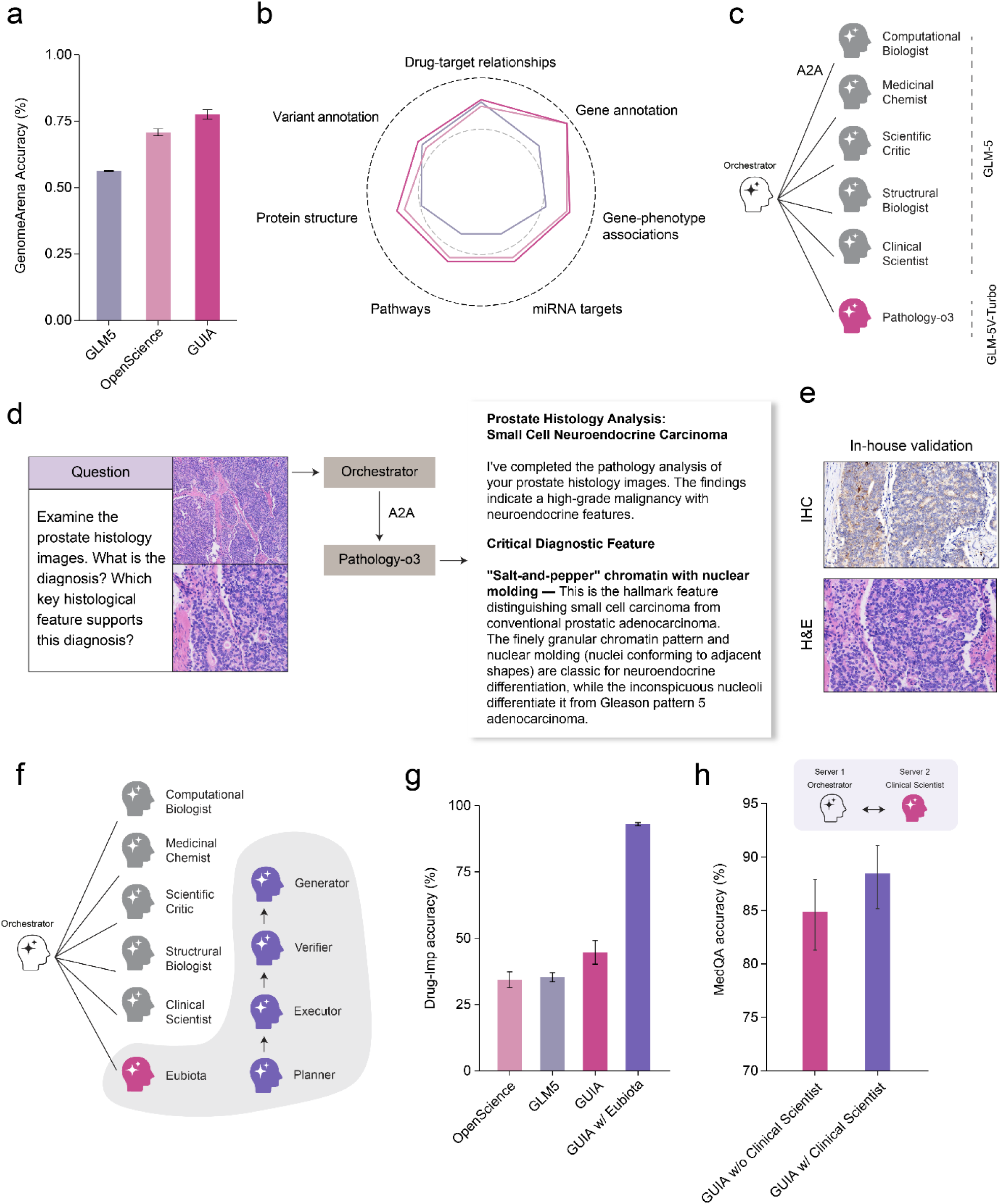
Evaluation of GUIA’s baseline performance and scalability. **a.** Average accuracy (*n* = 3) across 720 GenomeArena tasks. Error bars indicate the standard error of the mean (s.e.m). **b.** Per domain accuracy (*n* = 3). Biomedical tasks are split into seven categories: drug-target relationships, gene annotation, gene-phenotype associations, miRNA targets, pathways, protein structure, and variant annotation. **c.** GUIA collaborates with Pathology-o3, powered by GLM-5V-Turbo. **d.** Query and analysis of prostate histology images. **e.** Representative images of the prostate specimen stained with hematoxylin and eosin (H&E) and SYP. **f.** GUIA collaborates with Eubiota, an external multi-agent system consisting of four specialized agents: Planner, Executor, Verifier, and Generator. **g.** Average accuracy (*n* = 3) across 100 Drug-Imp tasks. Error bars indicate the s.e.m. **h.** Average accuracy (*n* = 3) across 150 MedQA tasks. The schematic depicts cross-server coordination with the Clinical Scientist agent. Error bars indicate the 95% confidence interval (CI).

As a second evaluation, we focused on a key limitation of GUIA’s in-house agents, namely their inability to analyze visual biomedical data owing to both the lack of relevant tools and the incompatibility of the underlying LLM. In fact, visual data represents a critical source of biomedical information, enabling disease diagnosis and the mapping of molecular features in-situ^19^. The importance of biomedical imaging has thus driven the rapid development of agentic systems designed to address various computational pathology tasks, although they remain largely isolated^9,20,21^. To demonstrate the flexibility of GUIA, we integrated a repurposed Pathology-o3 agent from Pathology-CoT^9^ into the agent network, thereby extending the platform’s capabilities to include WSI analysis (Fig. 3c). Of note, this integration was achieved through loose coupling without requiring tight system-level integration. This enabled Pathology-o3, powered by a distinct vision-language model (VLM), to collaborate seamlessly with GUIA’s in-house agents, highlighting the extensibility of our research-agent network (Fig. 3c).

We next sought to rigorously evaluate the performance of this extended vision-based capability. As this functionality is unavailable to agents powered by text-only LLMs, we focused on a real-world application to evaluate the robustness of GUIA’s results. Towards this, GUIA was presented with histology images from a clinical prostate specimen and tasked with proposing a diagnosis based on its histopathological features. Upon assessing the task, the Orchestrator delegated the analysis to the integrated Pathology-o3 agent (Fig. 3d). Interestingly, the tissue was diagnosed as small-cell neuroendocrine carcinoma of the prostate, a rare and aggressive subtype accounting for less than 1% of prostate cancer (PCa) cases^22,23^ (Fig. 3d). This prediction was compelling given the rarity of the disease and demonstrated that Pathology-o3’s interpretation was driven by definitive histological features rather than a bias toward more common PCa subtypes. To validate the agent’s diagnosis, we performed immunohistochemical (IHC) analyses, which revealed positive synaptophysin (SYP) staining, thus confirming neuroendocrine differentiation in the prostate tissue^24,25^ (Fig. 3e). Collectively, these results demonstrated the practical value of extending GUIA’s agent network through the loose coupling of third-party agents, enabling complementary expertise to be combined.

As a third evaluation, we explored whether GUIA could scale beyond single-agent integration, by collaborating with an external multi-agent platform on shared scientific tasks. To this end, Eubiota^11^, an agentic system for gut microbiome research was integrated into GUIA (Fig. 3f). Eubiota consists of four specialized agents dedicated to research planning, tool execution, evidence verification, and grounded knowledge synthesis. These agents leverage microbiome-specific tools, databases, and reasoning strategies to support the analysis of microbial data^11^. We therefore evaluated whether coupling GUIA with this third-party multi-agent framework resulted in improved performance on microbiome-oriented research tasks. Across tasks related to drug impacts on microbial growth (Drug-Imp), GUIA achieved an average accuracy of 93% following integration with Eubiota (GLM-5), substantially outperforming both standalone GUIA (GLM-5; 45%), the base LLM (GLM-5; 35%), and Open Science (GLM-5; 34%) (Fig. 3g).

For the fourth evaluation, we assessed whether cross-server collaboration could serve as a means of coordinating agent capabilities. To study this, we examined GUIA’s performance on medical tasks with and without access to the Clinical Scientist, an agent equipped with medical-specific tools, which was deployed on a geographically separate server. We showed that the collaboration with the Clinical Scientist resulted in a subtle improvement in MedQA^26^ task accuracy (Fig. 3h). Of note, although standalone GUIA already demonstrated strong medical performance, the collaboration with the Clinical Scientist yielded further improvements. This result demonstrates that a research-agent network can be beneficial even across distributed infrastructures, establishing cross-server agent collaboration as a paradigm for scientific research.

Collectively, these studies demonstrate that scientific capabilities can be expanded through the integration of specialized agents with non-overlapping expertise. While Pathology-o3 extended GUIA to support computational pathology, Eubiota introduced microbiome-specific reasoning and tool use, and Clinical Scientist contributed specialized medical expertise through cross-server collaboration. These findings suggest that research-agent networks can accumulate capabilities through interoperability, enabling scientific functionality beyond that of any individual constituent system.

### GUIA designs and executes a customized pipeline for drug discovery

To assess GUIA’s performance in real-world settings, we challenged the platform with three clinically unmet research problems that required cross-disciplinary expertise and multi-step scientific reasoning.

We first evaluated GUIA’s ability to propose and execute novel scientific solutions by applying it to in-silico drug discovery. Current computer-aided drug discovery (CADD) pipelines often rely on brute-force virtual screening, in which compound-protein interactions (CPIs) are evaluated across large chemical libraries to identify potential binders^27^. While powerful, this strategy is compute-intensive, time-consuming, and frequently associated with low hit rates. We therefore investigated whether GUIA could leverage scientific reasoning and domain-specific expertise to identify more efficient and innovative drug discovery strategies.

For demonstration, we selected KCNK9 as the drug target of interest, due to its oncogenic roles across multiple cancer types and the potential clinical significance of this finding^28,29^. We queried GUIA to design and execute an in-silico drug discovery pipeline that maximizes both success rate and operational efficiency (Extended Data Fig. 1a). The proposed pipeline was characterized by 5 distinct phases requiring expertise across multiple scientific domains. These include: (1) reference compound identification, (2) druggability assessment of chemical library, (3) similarity analysis, (4) target structure analysis, and (5) virtual screening for candidate prioritization (Extended Data Fig. 1b). Notably, while each of these components has been applied independently in various drug discovery settings, the proposed workflow with specific sequence in which the tools were deployed is distinct from existing approaches^30^. This pipeline enables efficient exploration of large compound libraries while balancing between predictive performance and resource utilization.

Briefly, GUIA first identified available protein structures and known inhibitors of KCNK9 as references. Using the SPECs chemical library, GUIA autonomously evaluated the druggability of each compound based on a series of metrics including the Lipinski’s Rule of Five, Quantitative Estimate of Drug-likeness (QED), and ADMET (Absorption, Distribution, Metabolism, Excretion, and Toxicity) properties^31–33^ (Extended Data Fig. 1c). Only compounds that satisfied predefined threshold were filtered for further investigation (Extended Data Fig. 1c). Next, pairwise Tanimoto similarity^34^ scores were calculated between each compound against known KCNK9 inhibitors to select for structurally similar candidates (Extended Data Fig. 1d, e). Finally, GUIA autonomously retrieved the highest quality KCNK9 protein structure from the PDB, employed AutoSite^35^ to identify putative binding pockets, and performed molecular docking using AutoDock Vina^36^ to predict CPI affinities (Extended Data Fig. 1f; Supplementary Fig. 2). This workflow substantially reduced the search space and prioritized chemically tractable and biologically relevant candidates, resulting in the identification of 19 novel drug hits (Extended Data Fig. 1g, h).

### GUIA proposes a novel spatial subtype classification for non-small cell lung cancer

We next evaluated GUIA’s capacity to reproduce findings from prior studies and extend those results to generate new biological insights. Towards this, we focused on imaging mass cytometry (IMC), a high-dimensional spatial proteomics technology that enables comprehensive characterization of cellular phenotypes and tissue organization^37–39^. Using an IMC dataset comprising 204 tumor histopathology images from 51 lung adenocarcinoma (LUAD) and 51 lung squamous cell carcinoma (LUSC) patients^39^, GUIA was tasked with reproducing established analyses and identifying novel patterns beyond those reported in the original study (Supplementary Fig. 3).

Given the cell type annotations derived from segmentation masks, GUIA recapitulated the distinct cellular compositions between LUAD and LUSC specimens^39^. Specifically, LUAD tumors exhibited significantly greater immune cell infiltration, whereas LUSC tumors were enriched for cancer cells (Fig. 4a). These cellular landscapes were mapped at the spatial level using cell segmentation data of the two cancer types (Fig. 4b). Moving forward, GUIA compared the ten cellular neighborhoods (CNs) reported in the original study, confirming that the differences between LUAD and LUSC extend beyond individual cell populations to local spatial niches (Fig. 4c-e).

**Figure 4.**
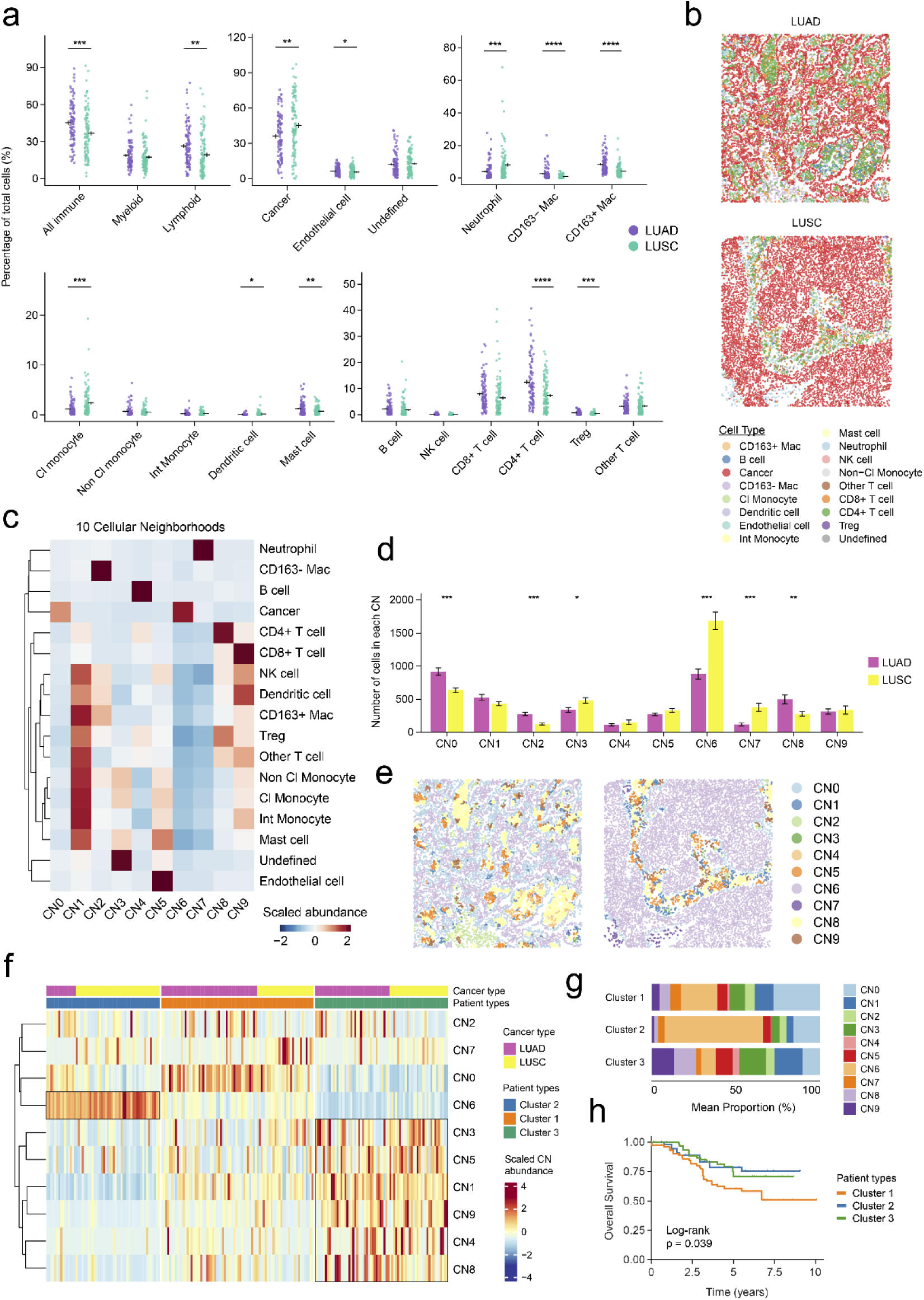
Analysis of IMC data from 102 NSCLC patients by GUIA. **a.** Prevalence of cell types across LUAD (*n* = 102) and LUSC (*n* = 102) samples. Statistical significance was assessed using two-tailed unpaired Student’s *t*-tests. Data are shown as mean ± s.e.m. **b.** Representative segmentation images of an LUAD and LUSC core, with cells coloured by cell types. **c.** Heatmap of cell type abundances across the 10 CNs. **d.** Abundance of each CN across LUAD and LUSC samples. Statistical significance was assessed using two-tailed unpaired Student’s *t*-tests. Error bars indicate the s.e.m. **e.** Representative segmentation images of an LUAD and LUSC core, with cells colored by CNs. **f.** Heatmap of NSCLC samples clustered by CN abundance. **g.** Prevalence of each CN across the three NSCLC spatial subtypes. **h.** Kaplan–Meier curve of overall survival with samples stratified by their NSCLC spatial subtype. Statistical significance was assessed using log-rank test. (\*\*\*\**p* < 0.0001; \*\*\**p* < 0.001; \*\**p* < 0.01; \**p* < 0.05).

Building upon these findings, we challenged GUIA to deduce novel biological patterns beyond those reported in the original study. GUIA reasoned that NSCLC tumors could be further stratified based on their CN composition, enabling a spatially defined subtyping system (Supplementary Fig. 3). Interestingly, while NSCLC subtypes have traditionally been identified by gene expression profiling^40,41^, GUIA proposed a spatially informed approach to tumor stratification. To test this hypothesis, GUIA autonomously performed K-means clustering of LUAD and LUSC samples based on their CN abundances, identifying three distinct NSCLC spatial subtypes (Fig. 4f). Cluster 1 exhibited a preference for spatial niches containing cancer cells, CD163-macrophages, and neutrophils; Cluster 2 was characterized by cancer cell enrichment and reduced immune infiltration, whereas Cluster 3 showed marked immune cell enrichment (Fig. 4g). Importantly, patients assigned to each clusters exhibited significantly different overall survival outcomes, demonstrating the prognostic value of the novel NSCLC classification framework (Fig. 4h).

### GUIA automates multi-omics analyses to uncover CASZ1 as a novel PCa biomarker

Thus far, we have demonstrated GUIA’s ability to tackle multidisciplinary research tasks and uncover previously unrecognized biological patterns. We next assessed its capacity to accomplish end-to-end research problems involving open-ended scientific reasoning.

We selected PCa biomarker discovery as the representative case study, with the objective of identifying novel AR coregulators that represent actionable vulnerabilities in PCa. This task has important clinical implications, as existing AR-targeted therapies provide only temporary clinical benefit before treatment resistance develops^42^. From a technical standpoint, this is an end-to-end research problem that encompasses multi-omics data analysis, clinical interpretation, literature-based knowledge discovery, pathological evaluation, and laboratory experimentation.

We constrained GUIA to analyze an in-house liquid chromatography–tandem mass spectrometry (LC–MS/MS) data generated from AR immunoprecipitation experiments across three PCa cell lines^43,44^. Using this dataset and the research objectives as input, GUIA autonomously designed and executed an end-to-end research workflow (Fig. 5a, b). The workflow began with the analysis of LC–MS/MS data to identify AR coregulators upregulated in one, two, or all three PCa cell lines (Fig. 5c). GUIA classified coregulators enriched in all three cell lines as high confidence and those enriched in two cell lines as moderate confidence. This was followed by the autonomous retrieval of PCa transcriptomic data from The Cancer

**Figure 5.**
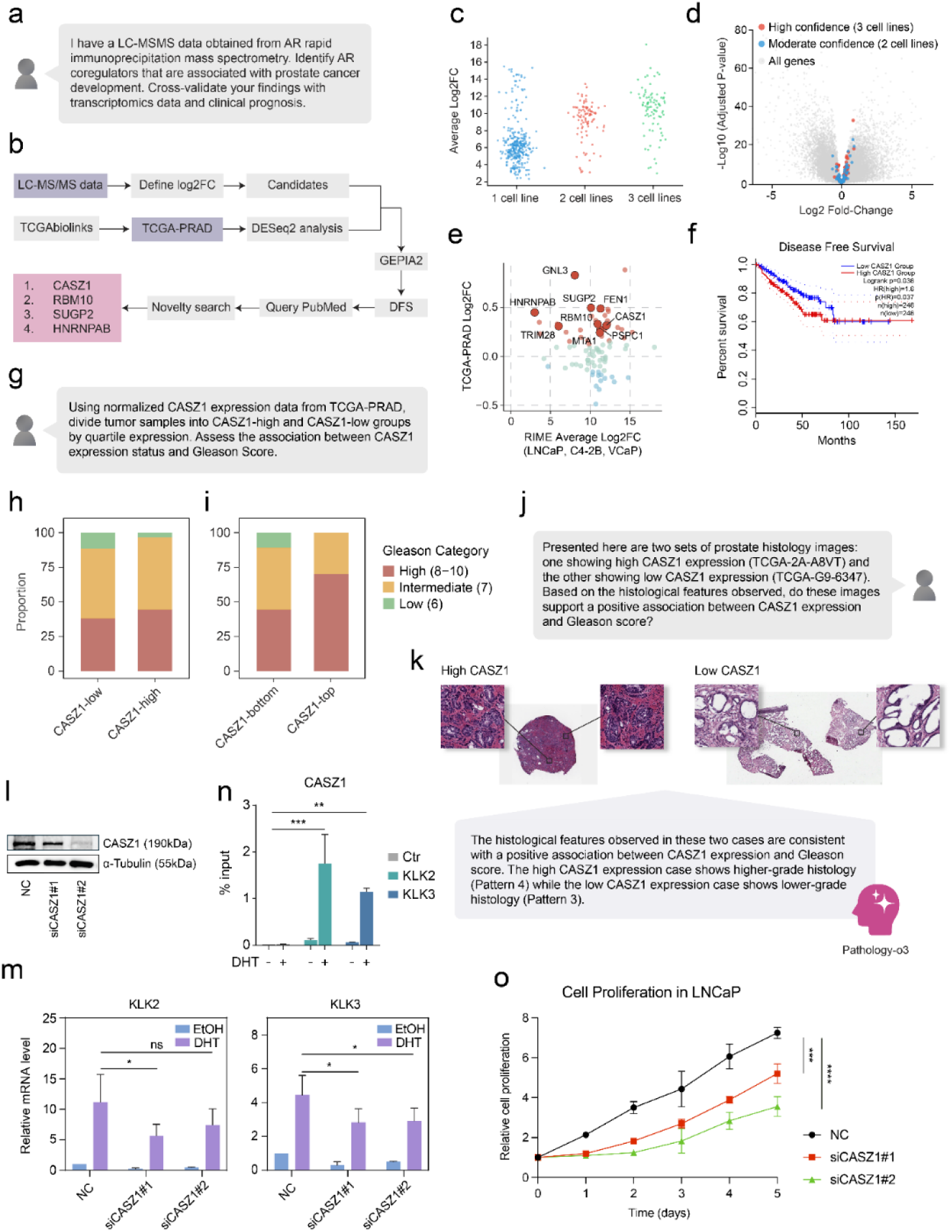
Identification of novel AR coregulators in PCa using GUIA. **a.** User query to initiate the case study. **b.** The multi-step workflow proposed by GUIA, involving multi-omics data analysis, clinical interpretation, and literature-based knowledge discovery. **c.** Upregulated AR coregulators identified by LC–MS/MS, enriched across one, two, or three PCa cell lines. **d.** Results from differential gene expression analysis of TCGA-PRAD data. Red dots indicate high-confidence coregulators identified in all three PCa cell lines by LC-MS/MS, whereas blue dots indicate coregulators identified in two PCa cell lines. **e.** Cross-comparison of log₂ fold changes in AR coregulators between LC-MS/MS and TCGA-PRAD. **f.** Kaplan–Meier curve of DFS stratified by median CASZ1 expression. The red line indicates high CASZ1 expression, and the blue line indicates low CASZ1 expression. **g.** User query for the correlation analysis of CASZ1 expression and Gleason score. **h.** Comparison of Gleason score distributions between CASZ1-high and CASZ1-low PCa samples. **i.** Comparison of Gleason score distributions restricted to the ten PCa samples with the highest and lowest CASZ1 expression. **j.** User query for the histopathological assessment of CASZ1 expression and Gleason score. **k.** WSI analysis of PCa samples with high and low CASZ1 expression. **l.** Western blot analysis of LNCaP cells treated with siCASZ1#1, siCASZ#2, and negative control (NC). **m.** Reverse transcription quantitative polymerase chain reaction (RT-qPCR) analysis of KLK2 and KLK3 expression in LNCaP cells treated with siCASZ1#1, siCASZ#2, and NC. Statistical significance was assessed using two-way analysis of variance (ANOVA). Error bars indicate s.e.m. **n.** CASZ1 ChIP-qPCR at AR target genes (KLK2 and KLK3) in LNCaP cells with or without DHT treatment (2 hours). Statistical significance was assessed using two-way ANOVA. Error bars indicate standard deviation (s.d). **o.** Cell proliferation in LNCaP cells following CASZ1 knockdown. Statistical significance was assessed using two-way ANOVA. Error bars indicate s.d. (\*\*\*\**p* < 0.001; \*\*\**p* < 0.001; \*\**p* < 0.01; \**p* < 0.05; ns, not significant at the 0.05 level).

Genome Atlas (TCGA-PRAD)^45^, which was subjected to differential gene expression analysis between tumor and normal prostate tissues (Fig. 5d). GUIA filtered for AR coregulators which were upregulated in both the proteomic and transcriptomic analyses and further queried GEPIA2^46^ to evaluate their association with patient disease-free survival (DFS) (Fig. 5e, f). Finally, leveraging access to the PubMed database, GUIA identified four novel AR coregulators, with CASZ1 being the highest-priority biomarker based on the consistency of supporting evidence.

Given the time-consuming and resource-intensive nature of biomarker validation, large-scale investigation into all candidates was not feasible. We therefore selected CASZ1 for further investigation. We first used GUIA to explore whether CASZ1 is associated with PCa progression, focusing on Gleason score as a measure of disease aggressiveness (Fig. 5g). GUIA autonomously stratified tumors into CASZ1-high and CASZ1-low groups based on upper and lower expression quartiles. Comparison of the two cohorts revealed a positive association between CASZ1 and Gleason score (Fig. 5h). This association was even more pronounced when the analysis was restricted to the ten PCa tumors with the highest and lowest CASZ1 expression (Fig. 5i). To verify this finding, we presented GUIA with histology images from matched patients and requested an unbiased assessment of their histopathological features (Fig. 5j). Leveraging GUIA’s collaboration with the Pathology-o3 agent, it concluded that the CASZ1-high tumor exhibited greater architectural distortion compared to the CASZ1-low tumor (Supplementary Fig. 4). Consistently, the predicted Gleason score was also greater in the CASZ1-high specimen, supporting an association between its expression and PCa progression (Fig. 5k).

To experimentally validate GUIA’s findings, we performed knockdown studies of CASZ1 in LNCaP cells (Fig. 5l). We showed that the silencing of CASZ1 significantly suppressed the expression of AR-regulated genes, indicating a requirement for CASZ1 in AR-mediated transcriptional activity (Fig. 5m). Furthermore, chromatin immunoprecipitation quantitative polymerase chain reaction (ChIP-qPCR) demonstrated the recruitment of CASZ1 to AR-binding sites associated with these target genes, suggesting that CASZ1 directly participates in AR-dependent transcriptional regulation (Fig. 5n). In line with its role in AR-mediated transcription, CASZ1 knockdown significantly impaired PCa cell proliferation, highlighting its contribution to tumor development (Fig. 5o). Collectively, these results establish CASZ1 as a novel AR coregulator that facilitates AR-dependent transcription in PCa cells.

### GUIA unlocks the collective potential of specialized scientific agents

In summary, we have demonstrated that interoperable research-agent networks can expand scientific capabilities through the integration of specialized agents with complementary expertise. Building on this architecture, we expanded the agent network to include 10 state-of-the-art (SOTA) scientific agents from both in-house and third-party sources. These agents include a Computational Biologist, Medicinal Chemist, Structural Biologist, Clinical Scientist, ChemCrow, CRISPR-GPT, Eubiota, GeneAgent, Pathology-o3, and Scientific Critic, each contributing domain-specific expertise across diverse areas of biomedical research^9–13^ (Fig. 6a). Collectively, this agent ecosystem enables GUIA to address a broad spectrum of scientific tasks, ranging from molecular and cellular biology to drug discovery, precision medicine, bioinformatics, pathology, organic chemistry, material science, gene-editing, and microbiology (Fig. 6b). Of note, while numerous biomedical agents have emerged, we prioritized the integration of agents with non-overlapping expertise to ensure stable task allocation and coordination. Looking ahead, this research-agent network can be further expanded by incorporating emerging scientific agents. Importantly, this extensible architecture positions GUIA as an interoperable research-agent network for scientific research.

**Figure 6.**
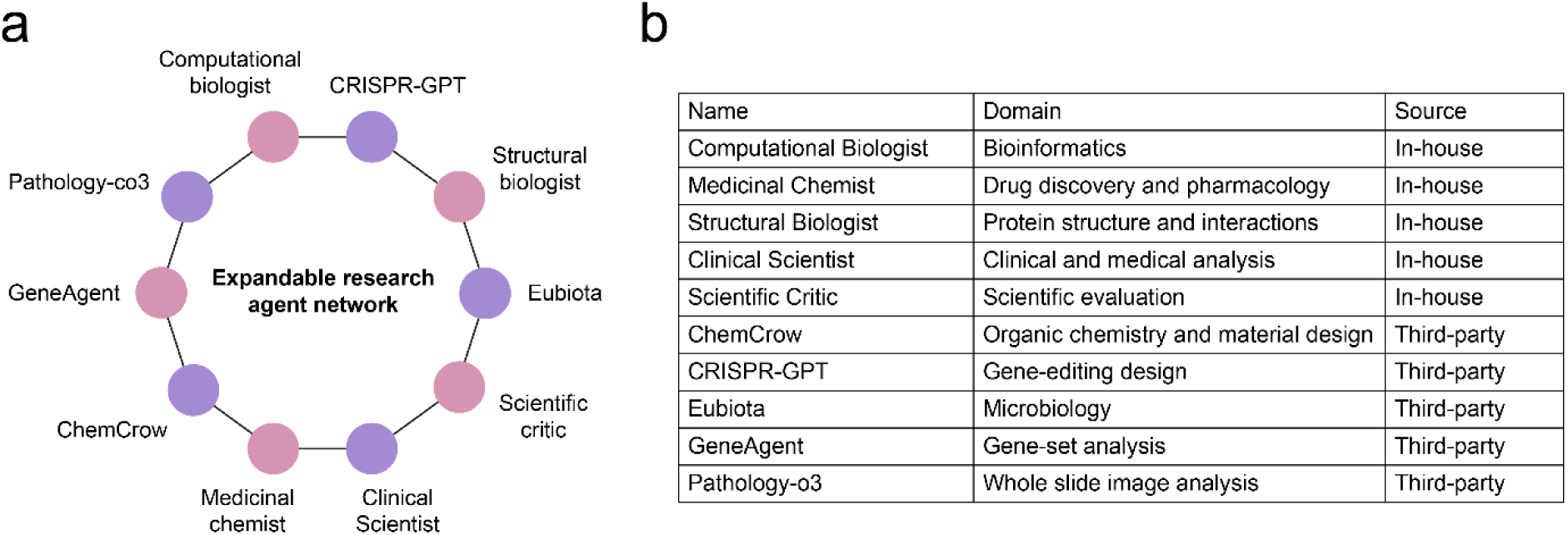
GUIA’s research agent network. **a.** An expandable research agent network comprising 10 specialized in-house and third-party agents. **b.** Description of the expertise associated with each specialized agent and the source from which it was derived.

## Discussion

In this study, we present GUIA, an interoperable research-agent network designed to facilitate seamless collaboration among specialized scientific agents. By integrating both in-house and third-party agents within a shared research ecosystem, GUIA enables scientific capabilities to accumulate through the coordinated use of complementary expertise across diverse research domains.

Previous advancements in agentic systems have largely focused on developing standalone agents optimized for specific research domains and workflows^9–13^. While these systems often demonstrate impressive performance within their specific areas of expertise, their operational isolation has created a fragmented landscape of scientific capabilities. This fragmentation limits the adoption of individual systems, as their expertise is confined to specific stages of larger scientific workflows. GUIA addresses this critical issue by establishing a unified research-agent network in which scientific agents from different platforms can interoperate within shared workflows. By coordinating complementary expertise across specialized agents, GUIA enables scientific capabilities to accumulate through collaboration, allowing complex research tasks to be addressed in ways that would be difficult for any individual agent to achieve alone. Furthermore, we show that such collaboration can be beneficial even across server boundaries, enabling agents hosted on different infrastructures to work together on shared scientific tasks.

GUIA offers significant advantages across four fronts. First, it removes the need for individual systems to be universally capable, enabling developers to focus on highly specialized expertise. Second, GUIA unifies the currently fragmented landscape of scientific agents, facilitating their coordination and adoption across interdisciplinary workflows. Third, GUIA overcomes infrastructure constraints by enabling agents hosted across different servers and computational environments to collaborate. This is increasingly important as specialized agents require unique hardware resources that are impractical to consolidate within a single infrastructure. Fourth, GUIA allows researchers to focus on answering scientific questions rather than the technical challenges of manually traversing between agent systems. By leveraging a collaborative research-agent network, GUIA lowers the technical barriers to conducting complex multidisciplinary research.

Despite the advantages of GUIA, several limitations remain. In its current implementation, file sharing is supported only among agents hosted on the same server. Cross-server collaboration is instead restricted to the exchange of structured messages. However, this limitation could be addressed through enhancements to the A2A framework that enable secure cross-server file transfer and shared data management. In addition, the current landscape of biomedical agents remains skewed towards areas such as bioinformatics and computational pathology, leaving other disciplines underserved. This constrains the breadth of capabilities available to GUIA. However, as the field matures and more specialized agents are released, GUIA’s expertise is expected to expand accordingly.

Like other agentic systems built upon LLMs, GUIA remains susceptible to reasoning and analytical errors. These may manifest as incorrect scientific interpretations and the generation of inaccurate results during research tasks. While GUIA incorporates multiple HITL mechanisms and provides opportunities for cross-agent verification, such safeguards cannot fully eliminate errors. The incorporation of specialized tools and data retrieval mechanisms may further alleviate these risks by grounding agent responses in authoritative resources. However, researchers should work with GUIA to verify key findings and ensure the accuracy of results.

In conclusion, GUIA serves as an interoperable research-agent network for scalable agent collaboration in scientific research. GUIA points toward a future in which scientific discovery is driven not by isolated agent systems, but by a cohesive network of agents working together on shared scientific tasks.

## Methods

### Configuring in-house agents

Specialized in-house agents are built using Google’s Agent Development Kit (ADK)^47^. ADK provides a flexible and robust environment for agent configuration, tool implementation, and session management. Each agent is configured through a system prompt comprising three components: (1) Title, which identifies the agent; (2) Role, which specifies its responsibilities within the agent network; and (3) Expertise, which determines the reasoning strategies, decision-making principles, and analytical approaches governing the agent’s actions. We find it useful to provide relaxed instruction prompts during agent configuration, as this maximizes their flexibility when handling a wide variety of tasks. Furthermore, we curated a collection of 97 tools across our in-house agents to support genomics, transcriptomics, proteomics, single-cell biology, spatial multi-omics, epigenomics, drug discovery, and other research domains. In addition, literature and information retrieval tools, such as PubMed and OpenAlex^48^ were provided to all in-house agents to facilitate knowledge discovery. These tools primarily operate within Bash, Python, and R.

Specialized agents are coordinated by a central agent known as the Orchestrator. The Orchestrator’s primary objective is to guide the overall research process by decomposing complex projects into manageable and parallelizable sub-tasks. These sub-tasks are delegated to the most relevant agents for execution.

### Harness engineering of in-house agents

Each in-house agent’s context is continuously monitored to prevent context overload. We employed a two-step approach for the effective management of the context window. First, existing context is summarized to distil key information, results, and task-relevant data generated by the agent. Subsequently, the context window becomes reinitiated with the distilled summary, thereby reducing context volume while preserving important information for future reasoning and task follow-up. To prevent context bloat caused by lengthy outputs from individual events, we developed a dedicated summarization sub-agent that condenses event outputs while preserving key information and results before injecting them into specialized agents.

Furthermore, we employed a dynamic context loading mechanism, whereby workflow references, guidelines, and best-practice behaviours are passed to agents on demand. This approach enables agents to retrieve task-relevant information when needed while minimizing context bloat. Specifically, workflow references and guidelines are modularized by design, enabling agents to retrieve specific components on demand rather than loading an entire workflow into the context window.

Importantly, agents are equipped with both semantic and episodic memory. This enables the holdout of key findings and research progress, which are continuously updated throughout the research session. Agents can retrieve these memory records when relevant, allowing previously generated knowledge and artifacts to be reused across different stages of a workflow.

### Communication between agents

We adapted Google’s Agent-to-Agent (A2A) protocol to support communication and collaboration across a network of research agents^14^. This provides a standardized architecture whereby agents communicate over HTTPS, with JSON-RPC 2.0 (Remote Procedure Call) serving as the data exchange format^14^. Both in-house and third-party agents were exposed through HTTP endpoints, each equipped with a dedicated AgentCard that outlines the agent’s metadata^14^. This AgentCard includes basic information about the agent, such as its name, description, research capabilities, service endpoint URL, supported modalities or data types, and other requirements. This communication protocol facilitates seamless collaboration among agents operating within the same server as well as across multiple servers, eliminating the need for tight system integration.

### Integrating third-party scientific agents

The following sections outline the process of integrating third-party scientific agents within GUIA’s research-agent network.

#### Pathologist-o3

Pathologist-o3 was converted into a standalone A2A-compatible agent by wrapping its original vision workflow with the Python A2A Software Development Kit (SDK)^49^. The adapter resolves the active GUIA session, selects one explicitly named pathology case, stages its thumbnail and pre-extracted ROI images, and runs low-magnification overview analysis, individual ROI assessment and final synthesis before saving JSON and PDF reports under the session’s directory. Notably, the original Pathologist-o3 system prompt and output fields were specific to detecting colorectal cancer metastasis in lymph nodes. We generalized these instructions into reusable prompts that support a wide range of pathology-related queries, allowing the same thumbnail–ROI–summary workflow to analyze diverse tissue types and address varying diagnostic objectives. This general mode is now the default, while the original colorectal lymph-node workflow remains available as an optional specialized mode. With Pathology-o3 alone, the agent still requires pre-extracted ROIs as input and does not support autonomous ROI extraction directly from WSIs.

#### ChemCrow

ChemCrow^12^ was converted into a standalone A2A-compatible system by wrapping its existing LangChain agent executor with the Python A2A SDK^49^. The adapter resolves the session ID and expands relative file paths from GUIA for ChemCrow navigation. An AgentCard advertises its chemistry capabilities, while incoming tasks are passed to the chemistry reasoning and tool-use workflow and returned to GUIA through A2A.

#### GeneAgent

GeneAgent’s gene-set interpretation and verification pipeline was wrapped in a standalone A2A server^13^. The adapter parses gene symbols from natural-language or structured requests, invokes the original evidence-verification cascade, and returns the resulting biological interpretation through A2A.

#### Eubiota

Eubiota^11^ was made A2A-compatible by placing an A2A execution layer around its existing Planner–Executor–Verifier–Generator workflow. The adapter resolves GUIA sessions, supplies uploaded files, creates an isolated workspace, executes the original multi-turn reasoning loop, and saves responses as well as results into the corresponding session directories.

#### CRISPR-GPT

CRISPR-GPT Lite^10^ was exposed as a standalone A2A agent by wrapping its custom gene-editing state machine. The adapter forwards orchestrator requests to the CRISPR planning stage, preserves the original safety checks, and returns knockout, Cas-system, guide-RNA, and off-target guidance through A2A.

### Benchmarking GUIA across multiple scientific domains

The following sections outline the process of evaluating GUIA’s performance across multiple benchmarks. Default temperature settings and model parameters were used for all evaluations to ensure consistency.

#### GenomeArena

We evaluated GUIA’s performance on biomedical tasks using GenomeArena, a benchmark dataset comprising 720 biomedical-oriented multiple choice questions (MCQs)^18^. These tasks encompass a range of domains, including (1) drug-target relationships, (2) gene-phenotype association, (3) gene annotation, (4) miRNA targets, (5) pathway knowledges, (6) protein structures, and (7) variant annotation. The MCQs were originally organized into domain-specific partitions. Each partition was submitted to the corresponding platform for evaluation, and performance was assessed by calculating the accuracy based on the generated responses. To account for variability, each task was repeated three times, and the reported accuracy was averaged across the three runs.

#### Drug-Imp

GUIA’s extensibility was assessed through the benchmarking of Drug-Imp, a dataset consisting of 100 tasks related to drug impact on microbial growth^11^. Similar to GenomeArena, questions were partitioned into sets of 10 MCQs each and submitted to the corresponding platforms for analysis. Performance was then assessed based on the average accuracy of generated responses across three separate runs.

#### MedQA

The effectiveness of cross-server collaboration was assessed using MedQA^26^. We randomly sampled 150 questions, partitioned them into three sets of 50 questions, and submitted these to each platform for analysis. Similarly, performance was evaluated based on the average accuracy of the generated responses across three independent runs.

### Workflow for identifying novel AR coregulators

Given the in-house LC-MS/MS data and user prompt as inputs, GUIA designed a computational workflow comprising four critical phases: (1) analysis of LC-MS/MS data to identify high-confidence AR coregulators across three PCa cell lines; (2) validation of the proteomic findings using TCGA-PRAD transcriptomic data; (3) disease-free survival (DFS) analysis using TCGA-PRAD data through GEPIA2; and (4) generation of actionable recommendations informed by literature evidence. For each phase, the Orchestrator delegated the task to the most relevant agent for execution. High-confidence AR coregulators were identified based on their log_2_ fold-change values across the PCa cell lines. TCGA-PRAD data were autonomously downloaded by GUIA using TCGAbiolinks^50^, followed by differential gene expression analysis with DESeq2^51^. GUIA further leveraged its access to GEPIA2^46^ to obtain DFS data for the selected candidates. Finally, novelty assessments were conducted through REST-API access to PubMed.

### Workflow for in-silico drug discovery

GUIA designed a computational workflow comprising five critical phases: (1) collection of PDB structures and existing KCNK9 inhibitor information; (2) design of an optimized KCNK9 compound library by filtering compounds based on drug-like properties; (3) prioritizing compounds with chemical structures similar to those of existing inhibitors; (4) molecular docking against KCNK9 structures; (5) candidate prioritization.

Available KCNK9 protein structures and existing KCNK9 inhibitors were autonomously retrieved by GUIA from the PDB, PubChem, and ChEMBL, respectively^15,52,53^. Compounds from the SPECs chemical library were evaluated for their drug-likeness using multiple criteria, including Lipinski’s Rule of Five, the Quantitative Estimate of Drug-likeness (QED), and ADMET (Absorption, Distribution, Metabolism, Excretion, and Toxicity) properties^31–33^. GUIA subsequently computed pairwise Tanimoto similarity scores between the filtered compounds and known KCNK9 inhibitors. The top 1,000 most similar compounds for each reference inhibitor were selected for further evaluation. Putative ligand-binding pockets were identified using AutoSite, and molecular docking simulations were performed with AutoDock Vina to predict CPI affinities. Compounds were then ranked according to their predicted affinities, and GUIA applied a cutoff of less than −10 kcal/mol to identify 19 candidate compounds.

### Analysis of IMC data from 102 NSCLC patients

GUIA autonomously re-processed an IMC data to reproduce various components from the original findings^39^. First, using cell type annotations derived from segmentation masks, GUIA calculated the proportion of each cell type relative to the total number of cells in each sample. Differences in cellular composition between LUAD and LUSC were subsequently evaluated. CN identities, which were derived in the original study from each cell’s ten nearest neighbors, were provided as input. GUIA autonomously reproduced the published CN-based analyses to verify and validate the original findings.

GUIA proposed multiple analyses to uncover novel biological patterns within the IMC data. One of the highest-priority suggestions was patient CN signature clustering, in which tumors were stratified according to their CN composition profiles. GUIA hypothesized that variations in spatial organization could serve as the basis for a novel tumor microenvironment subtyping system. Towards this, GUIA autonomously performed K-means clustering to identify three patient clusters characterized by distinct microenvironmental architectures. Furthermore, GUIA analyzed the cellular composition of each patient cluster and performed overall survival analyses to evaluate their clinical outcomes.

### IHC analysis

A prostate tissue specimen in the form of formalin-fixed, paraffin-embedded (FFPE) sections were sourced from Shanghai Outdo Biotech. IHC staining for SYP was performed on the FFPE tissue section. The section was deparaffinized in xylene and rehydrated through an ethanol gradient. Heat-induced antigen retrieval was then carried out using Target Retrieval Solution (Agilent Dako, S2367; pH 9.0). Following antigen retrieval, the section was incubated overnight at 4 °C with an anti-SYP primary antibody (Santa Cruz Biotechnology, sc-17750). After thorough washing with 1× PBS, the section was incubated with secondary antibody for 2 hours, and immunoreactivity was visualized using a DAB Detection System (BOSTER).

### Cell culture

The human PCa cell line LNCaP (ATCC CRL-1740; RRID: CVCL_1379) was obtained from the American Type Culture Collection (ATCC, Manassas, VA, USA). Cells with passage numbers below 20 were used for all experiments. Mycoplasma contamination was routinely assessed using the MycoAlert Mycoplasma Detection Kit (Lonza), and all cultures tested negative throughout the study. Cells were maintained in RPMI-1640 medium (Gibco) supplemented with 10% fetal bovine serum (FBS; Gibco) and 1% penicillin-streptomycin (Gibco). Cultures were incubated at 37°C in a humidified atmosphere containing 5% CO2.

### siRNA transfection

Cells were seeded in 6-well plates 24 hours prior to transfection and allowed to reach approximately 60% confluence. Dicer-substrate small interfering RNAs (DsiRNAs) were purchased from Integrated DNA Technologies (Supplementary Table. 1). For each well, siRNAs were diluted to a final concentration of 20 nM in 42.5 μL transfection buffer (GenePharma, China), followed by the addition of 15 μL siRNA-mate transfection reagent (GenePharma, China). After gentle mixing, the transfection complexes were added directly to the cells. Cells were maintained in complete growth medium supplemented with serum and antibiotics and incubated at 37°C in a humidified atmosphere containing 5% CO₂.

### RT-qPCR analysis

Total RNA was extracted using the RNeasy Mini Kit (QIAGEN, Cat. No. 74106) according to the manufacturer’s instructions. First-strand cDNA was synthesized using 4× All-in-One qRT Super Mix (Vazyme). RT-qPCR was performed with PerfectStart® Universal Green qPCR SuperMix (TransGen Biotechnology, Cat. No. AQ631), and relative gene expression levels were determined using the corresponding amplification signals. Experiments were performed in two biologically independent replicates (*n* = 2).

### Western Blotting

Cells were seeded at equal densities in 6-well plates containing 1.5 mL of complete medium and transfected with siRNAs for the indicated durations. Following treatment, cells were collected by scraping, resuspended in culture medium, and pelleted by centrifugation at 16,000 × g for 20 minutes. Total protein was extracted and quantified using the Bradford protein assay. For immunoblotting, 30 μg of total protein from each sample was mixed with 4× Laemmli sample buffer (Bio-Rad, #1610747) and denatured at 95°C for 5 minutes. Protein samples were separated by SDS-PAGE using a Bio-Rad electrophoresis system and transferred onto membranes at a constant current of 300 mA for 3 h. A pre-stained protein ladder (10-250 kDa, Epizyme, WJ103) was included as a molecular weight marker. Following transfer, membranes were washed with PBST and blocked with 5% bovine serum albumin (BSA) in PBST for 2 hours at room temperature. Membranes were then incubated overnight at 4°C with primary antibodies, including anti-CASZ1 (Santa Cruz Biotechnology, sc-398303). After washing with PBST, membranes were incubated with the appropriate HRP-conjugated secondary antibodies for 2 hours at room temperature. Protein bands were detected using an enhanced chemiluminescence (ECL) detection reagent (Epizyme) and visualized with a ChemiDoc™ imaging system (Bio-Rad). Experiments were performed in two biologically independent replicates (*n* = 2).

### CASZ1 ChIP-qPCR

ChIP-qPCR was performed as previously described^43,44,54^. Briefly, 1 × 10⁷ LNCaP cells were seeded in 15-cm dishes and cultured in hormone-deprived medium for 24 hours. Cells were treated with either ethanol (EtOH) vehicle control or 100 nM DHT for 2 hours before cross-linking with formaldehyde (Sigma, 252549). The cross-linked cells were scraped, collected, and incubated with an anti-CASZ1 antibody (Santa Cruz Biotechnology, sc-398303) that had been pre-coated onto Protein G Dynabeads (Invitrogen, 10009D). DNA was subjected to overnight precipitation followed by de-crosslinking at 65°C. The products were subjected to qPCR. Experiments were performed in three biologically independent replicates (*n* = 3).

### Cell Proliferation assay

LNCaP cells were seeded in 6-well plates and transfected with DsiCASZ1 or a negative control (NC) siRNA. After 24 hours of transfection, cells were harvested by trypsin and counted. Equal numbers of viable cells were subsequently seeded into 96-well plates for proliferation analysis. Cell proliferation was assessed daily using the AlamarBlue Cell Viability Reagent (Invitrogen, DAL1100) according to the manufacturer’s instructions. Fluorescence signals were measured at the indicated time points and used to evaluate cell proliferation. Experiments were performed in three biologically independent replicates (*n* = 3).

## Data availability

TCGA-PRAD WSI data are available from the GDC Data Portal at https://portal.gdc.cancer.gov/. Proteomics data from AR immunoprecipitation LC-MS/MS are available at the ProteomeXchange Consortium, under dataset ID PXD067138^43^. IMC data for NSCLC are available via Zenodo at https://doi.org/10.5281/zenodo.14625562 (ref.^39^). The GenomeArena (https://github.com/ksenia007/alvessa_agent) (ref.^18^) and Drug-Imp (https://github.com/lupantech/Eubiota) (ref.^11^) benchmarks are available from Github. The MedQA (https://huggingface.co/datasets/bigbio/med_qa) (ref.^26^) questions are available from Hugging Face.

## Code availability

The A2A protocol used for GUIA’s multi-agent collaboration framework is publicly available at https://github.com/a2aproject/A2A (ref.^14^). Third-party agents used for GUIA’s integration are available on GitHub at https://github.com/ur-whitelab/chemcrow-public (ref.^12^), https://github.com/lupantech/Eubiota (ref.^11^), https://github.com/ncbi-nlp/GeneAgent (ref.^13^), https://github.com/cong-lab/crispr-gpt-pub (ref.^10^), and https://github.com/zhihuanglab/Pathology-CoT (ref.^9^). To access GUIA, please complete the registration form at https://qz-l.com/ziePfa.

## Acknowledgements

This research was funded by grants provided by the University of Macau (MYRG2022-00204-FHS, MYRG-GRG2023-00189-FHS-UMDF, and MYRG-GRG2024-00178-FHS) and the Macau Science and Technology Development Fund (0137/2020/A3 and 0030/2025/AMJ). We thank the members of the laboratories led by Edwin Cheung, Wei Ge, and Xianting Ding for their valuable feedback and insightful discussions. The authors gratefully acknowledge the support and assistance provided by the Faculty of Health Sciences and the Information and Communication Technology Office at the University of Macau.

## Author information

### Authors and Affiliations

Faculty of Medicine, Faculty of Health Science, Ministry of Education Frontiers Science Center for Precision Oncology, University of Macau, China

Tin Long Cheong, Xinchen Ji, Ying Wang, Cuiting Zhang, Bo Li, Yue Zhou, Jiabao Huang, I Wu, Ao Li & Edwin Cheung

### Contributions

T.L.C. and E.C. conceptualized the study and contributed the methodology; T.L.C. developed GUIA with contributions from Y.Z. and J.H.; T.L.C, Y.Z., and Y.W. contributed to benchmark evaluations. X.J., C.Z., and B.L. contributed to laboratory experiments. T.L.C., I.W., and A.L., contributed to GUIA testing and assessment.

---

**Extended Data Figure 1.**
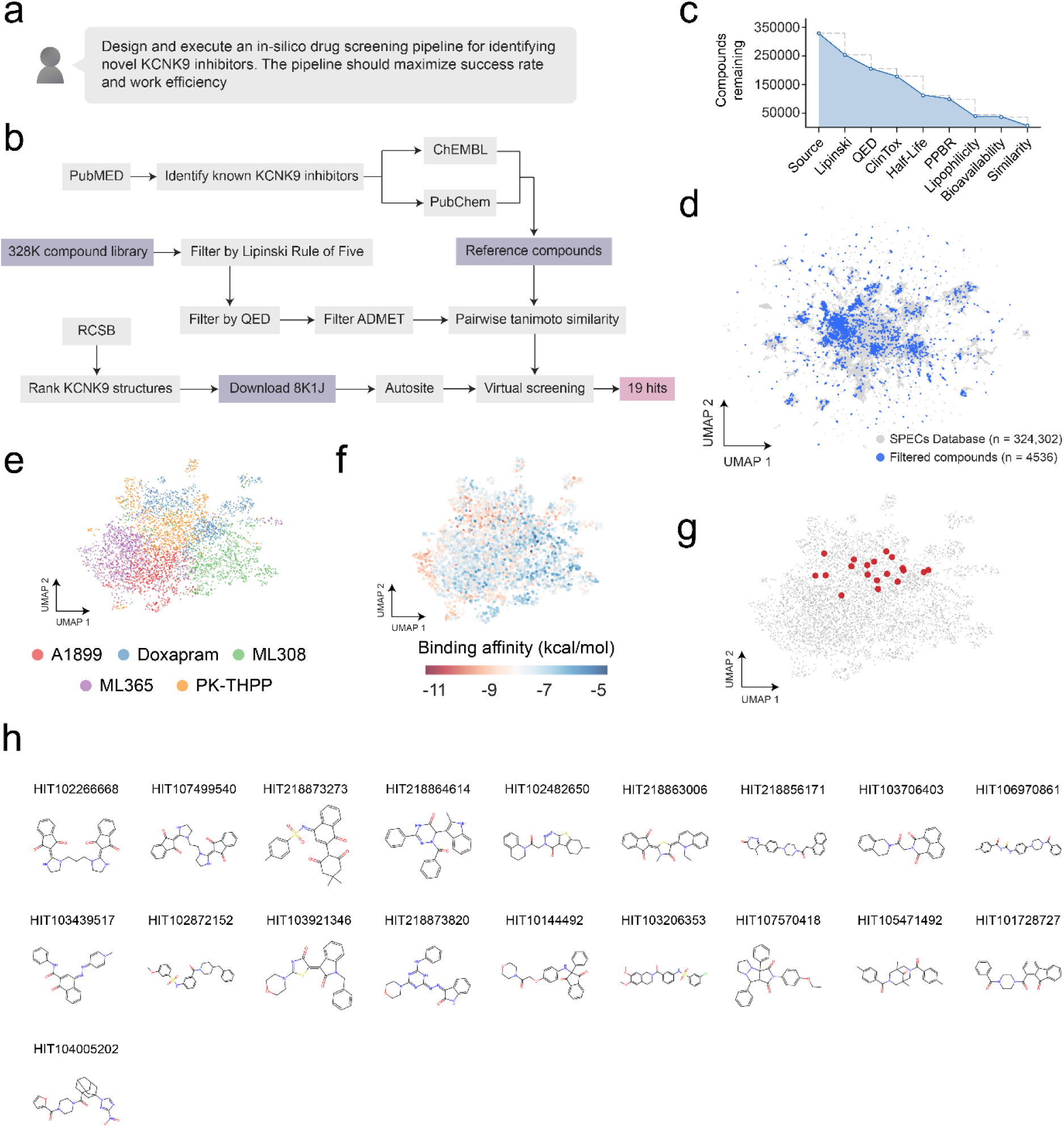
Identification of KCNK9 inhibitors using GUIA. **a.** User query to initiate the case study. **b.** Drug discovery workflow proposed by GUIA, involving the selection of reference KCNK9 inhibitors, assessment of compound druggability, compound similarity analysis, identification of putative ligand-binding pockets, and molecular docking. **c.** Chemical library size at each stage of the multi-step filtering process. **d.** Uniform Manifold Approximation and Projection (UMAP) visualization of over 324,000 compounds from the SPECs chemical library using Morgan fingerprints. Blue dots indicate compounds retained after GUIA filtering. UMAP of 4536 filtered compounds colored by **e.** the reference inhibitor they resemble, **f.** AutoDock Vina affinity scores, and **g.** the final 19 candidates. **h.** Chemical structures of the 19 drug candidates.

**Supplementary Figure 1.**
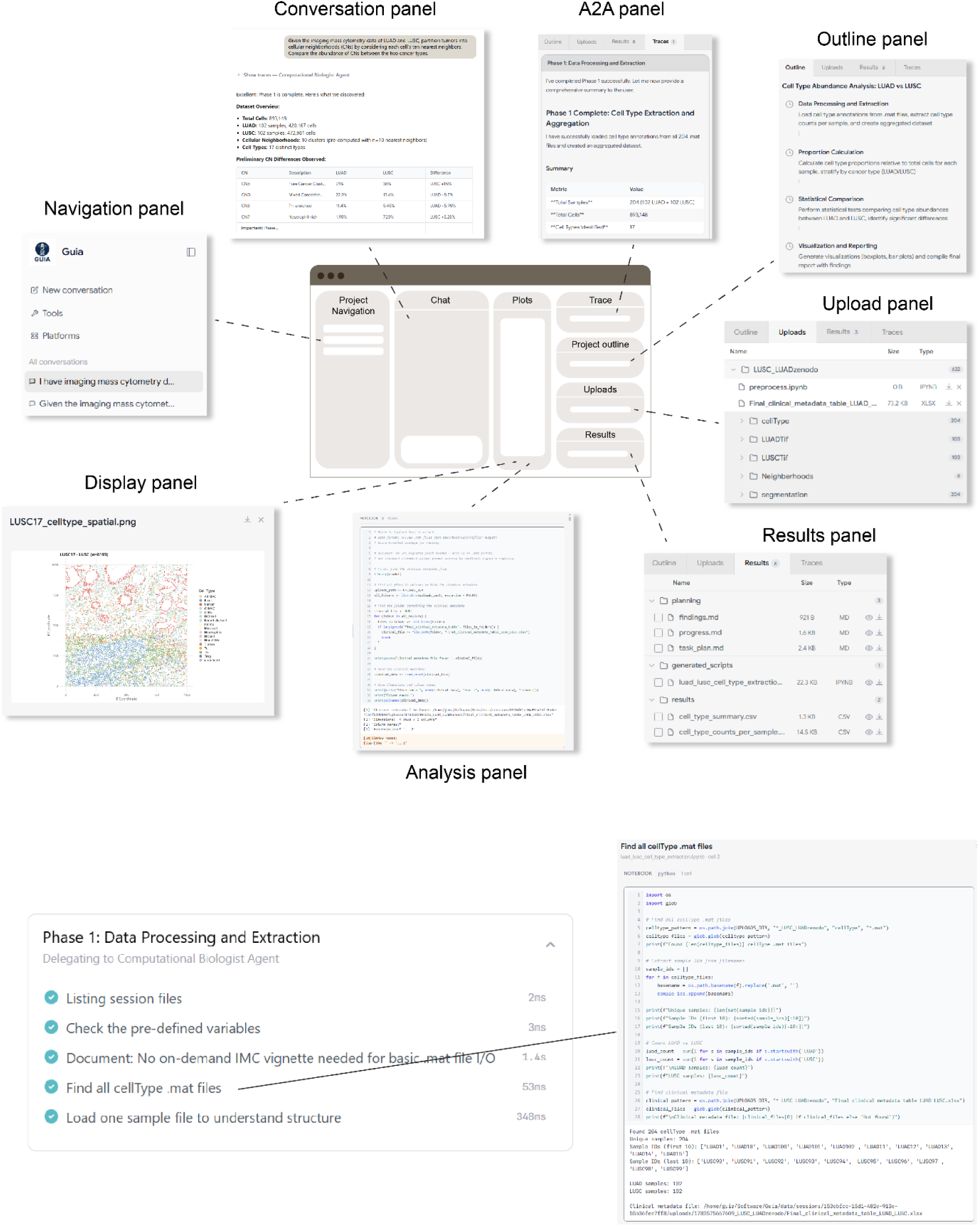
Frontend panel for real-time agent activity tracing

**Supplementary Figure 2.**
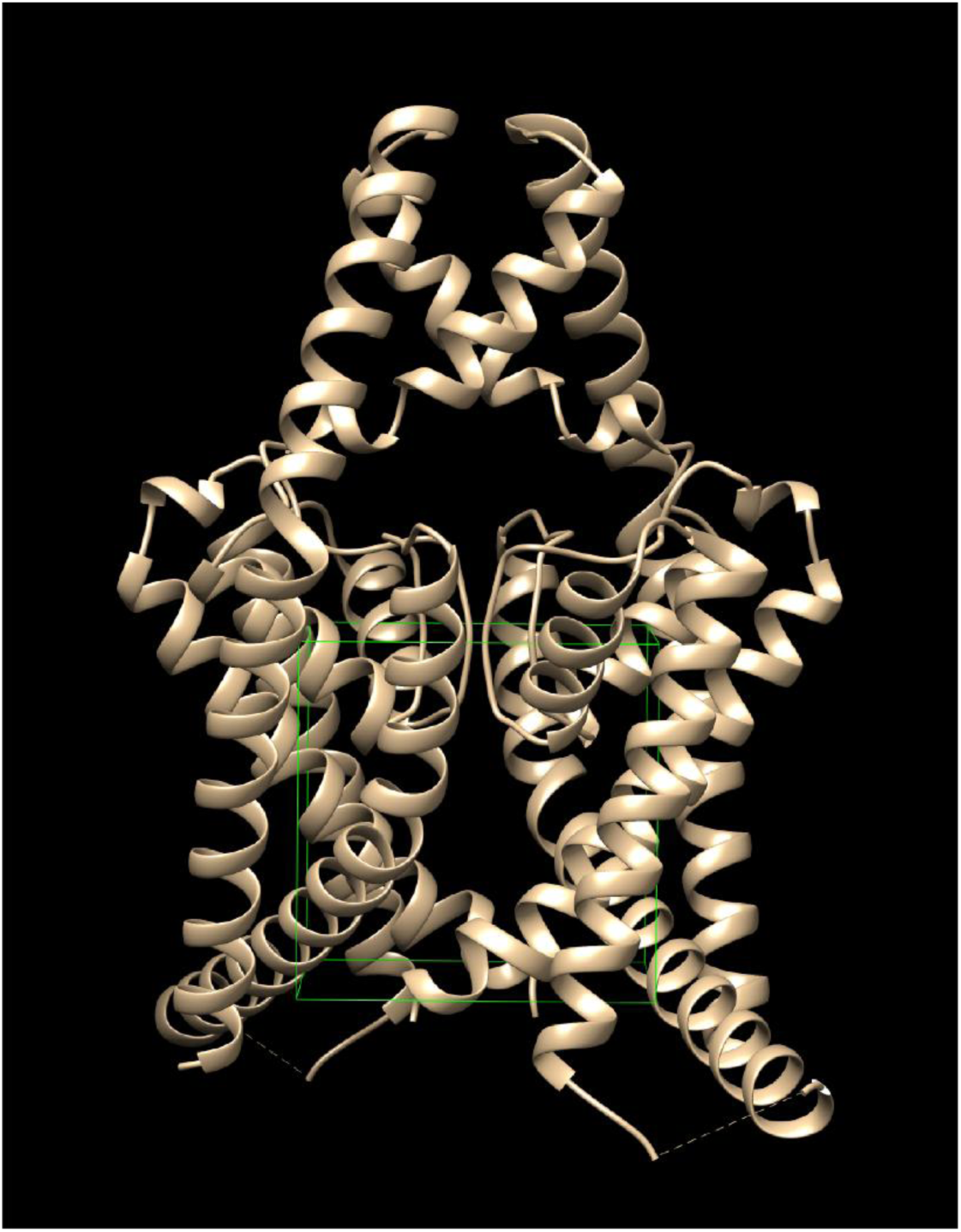
Highest-scoring ligand-binding site on KCNK9 predicted using AutoSite. The predicted site is consistent with the expected binding site.

**Supplementary Figure 3.**
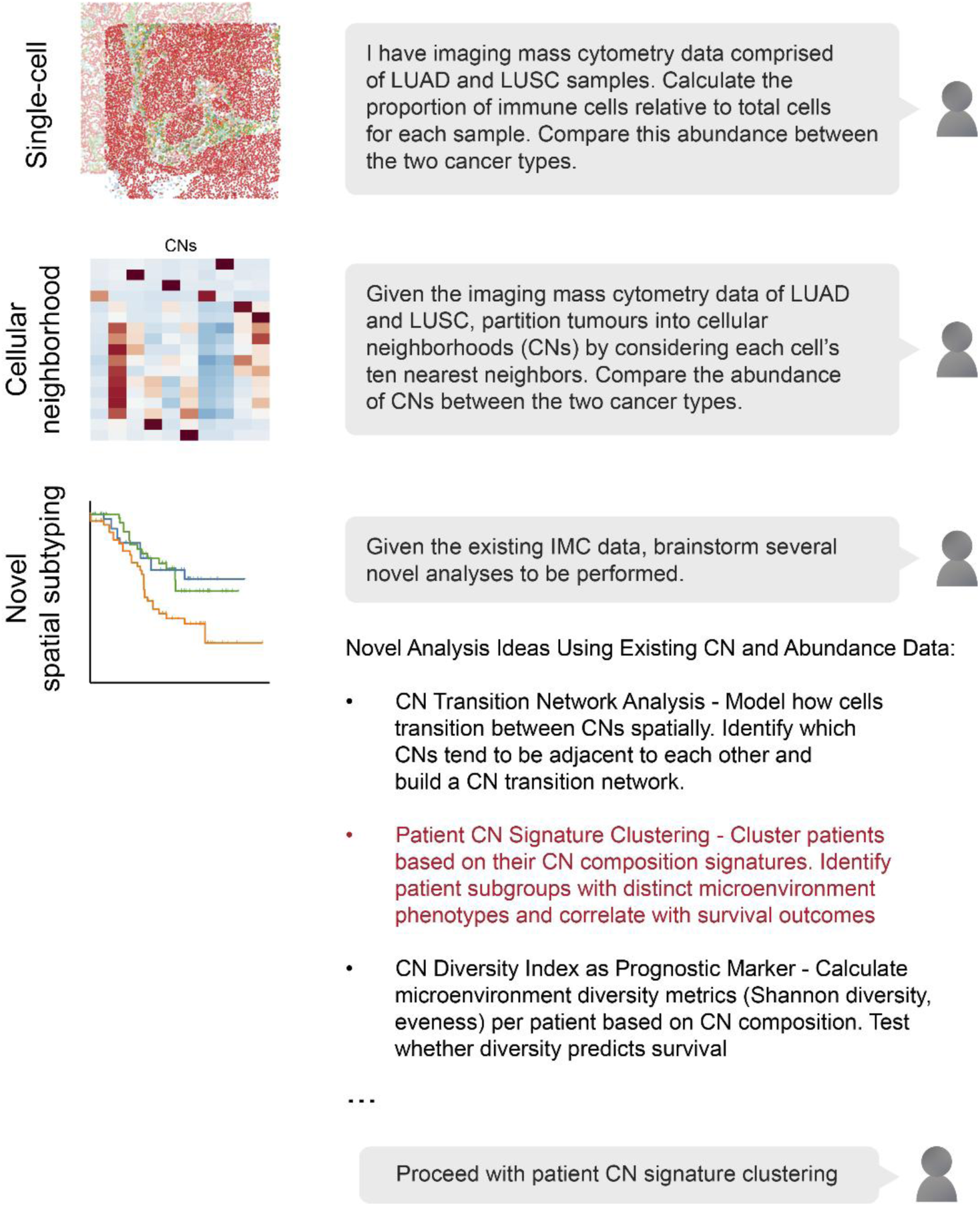
User queries corresponding to each step of IMC analysis.

**Supplementary Figure 4.**
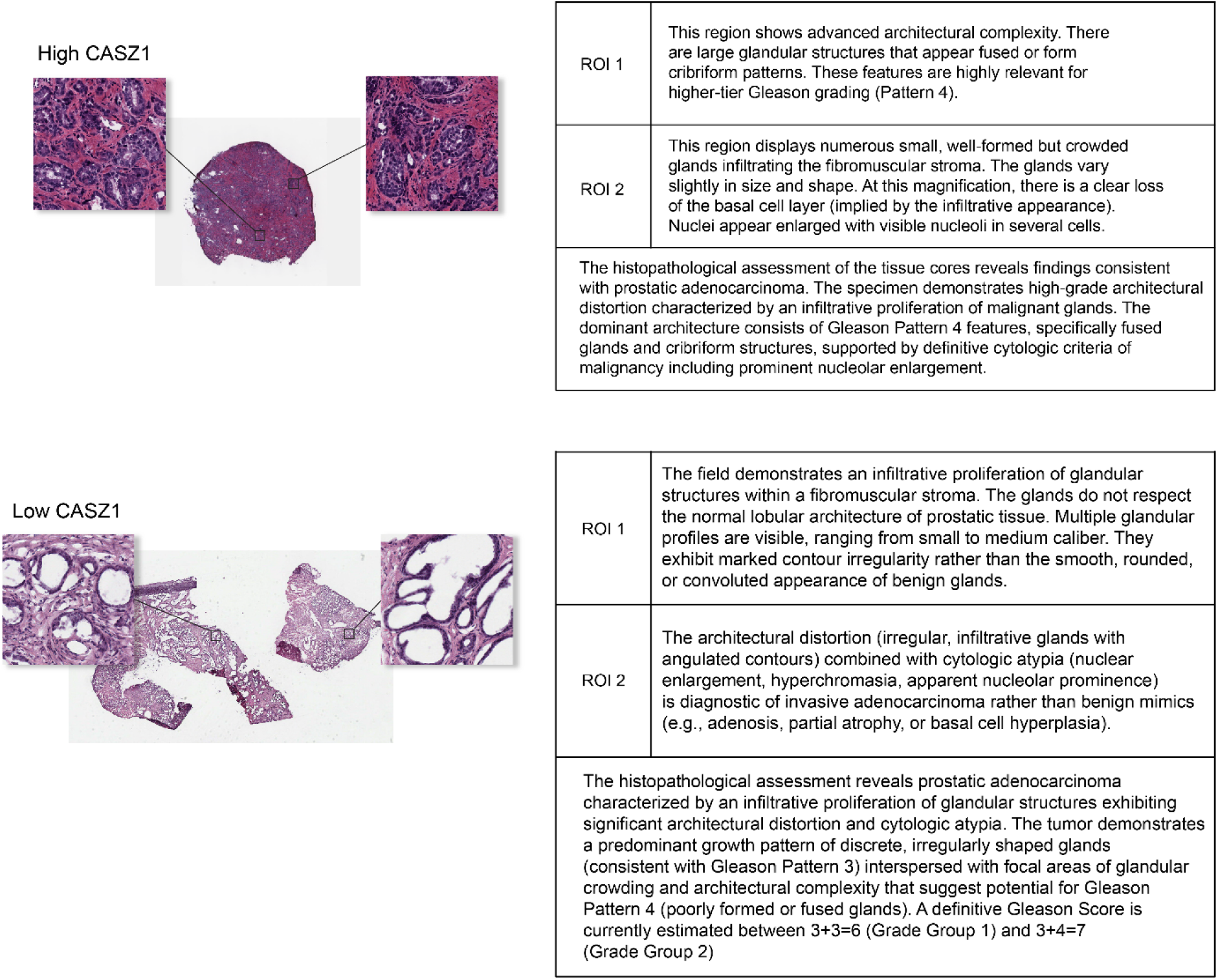
WSI analysis of CASZ1-high and CASZ1-low prostate tumors. A low-magnification thumbnail, and two high-magnification regions of interest (ROIs) were provided for each tumor.

**Supplementary Table 1.** Sequence information for DsiCASZ1.

| DsiCASZ1 sequences |  |
| --- | --- |
| DsiCASZ1.1 | DsiCASZ1.2 |
| 5'-<br>rCrArCrUrGrArArGrArUrGrUrArArArCrArUrUrUr<br>ArCrCAG-3' | 5'-<br>rGrUrArCrCrUrGrArArGrUrCrArArCrCrUrUrCrUr<br>CrCrAAA-3' |
| 5'-<br>rCrUrGrGrUrArArArUrGrUrUrUrArCrArUrCrUrUr<br>CrArGrUrGrArC-3' | 5'-<br>rUrUrUrGrGrArGrArArGrGrUrUrGrArCrUrUrCrAr<br>GrGrUrArCrUrC-3' |

